# Natural selection towards wild-type in composite cross populations of winter wheat

**DOI:** 10.1101/678938

**Authors:** Samuel Knapp, Thomas F. Döring, Hannah E. Jones, John Snape, Luzie U. Wingen, Martin S. Wolfe, Michelle Leverington-Waite, Simon Griffiths

## Abstract

Most of our crops are grown in monoculture with single genotypes grown over wide acreage. An alternative approach, where segregating populations are used as crops is an exciting possibility, but outcomes of natural selection upon this type of crop are not well understood. We tracked allelic frequency changes in evolving composite cross populations (CCPs) of wheat grown over ten generations under organic and conventional farming. At three generations, each population was genotyped with 19 SSR and 8 SNP markers. The latter were diagnostic for major functional genes. Gene diversity was constant at SSR markers but decreased over time for genic markers. Population differentiation between the four locations could not be detected, suggesting that organic vs. non-organic crop management did not drive allele frequency changes. However, we did see changes for genes controlling plant height and phenology in all populations independently and consistently. We interpret these changes as the result of a consistent natural selection towards wild-type. Independent selection for alleles that are associated with plant height suggests that competition for light was central, resulting in the predominance of stronger intraspecific competitors, and highlighting a potential trade-off between individual and population performance.

## Introduction

Successful crop production depends on varieties that are well adapted to a target environment (Cooper and Hammer, 1996; Atlin et al., 2017) but sufficiently widely adapted so that breeding and seed production is economically viable. As a result, a large proportion of the harvested area is occupied by a few major inbreeding crops (e.g. wheat, barley, rice) and within any one farm, large blocks of each crop comprise single genotypes. This bears risks of vulnerability to diseases (Brown and Hovmøller, 2002) and limited adaptability to local conditions (Mercer and Perales, 2010).

A potential response to these drawbacks is the use of genetically diverse populations instead of clonal crops (Litrico and Violle, 2015). Crop populations can be created by mixing different varieties (Finckh and Wolfe, 2006) or by inter-crossing varieties and mixing the progenies (Suneson, 1956), which, in combination with harvesting and re-sowing each generation, is called evolutionary plant breeding (Suneson, 1956; Döring et al., 2011). Genotypes better adapted to local conditions should have more progeny and thus increase in frequency and over time could result in better locally adapted genotypes and increased grain yield (Döring et al., 2011).

Early wheat studies describe yield increases and with reported rates of genetic gain comparable to those mainstream breeding (Suneson, 1956). Similar results were reported by Allard (1988) and for biotic stress, Le Boulc’h et al. (1994) found increased resistance to powdery mildew as did Paillard et al. (2000). Furthermore, it was found that diverse winter wheat populations across France showed a differentiation in phenological development (Rhoné et al., 2010). Populations grown over several generations in Northern France with colder winters flowered later than populations grown in Southern France with risk of drought at the end of the growing season. Recently, Bertholdsson et al. (2016) showed that seedling traits of winter wheat CCPs were differentially selected in organic vs. conventional management systems, with the populations maintained under organic management showing an increase in seedling root length and root weight, while populations under conventional management showed no such increase. This suggests that the selection CCPs are subjected to can lead to adaptation to locally prevailing conditions and management systems.

These positive results were achieved in spite of the trade-off between individual plant fitness and population performance (Weiner et al., 2010; Denison, 2012; Anten and Vermeulen, 2016). Natural selection acts on individuals but population performance is the central variable in crop production. Individual fitness in a population strongly depends on competition-related traits such as plant height, but investment by individual plants in competition may reduce their potential for grain yield (Weiner et al., 2010). Accordingly, harvest index in cereals, i.e. the proportion of grain yield in total biomass, decreases with increasing intra-specific completion among crop plants with increasing density (Weiner and Freckleton, 2010).However, under no-herbicide conditions of organic farming, where weeds are often more abundant than in conventional cropping systems (Gabriel et al., 2013), the same competition-related traits may be of advantage.

Our main objective was to find out whether selection can lead to genetic differentiation reflecting adaptation to different management conditions, and if the signature of this selection can be detected for a set of genes with particular importance for competition. To relate the function of selected alleles to these genetic signatures, we also evaluated allelic effects on plant height, heading date, yield and yield components in pure stands and on individual plants in the CCPs. We conducted this investigation on CCPs of winter wheat grown with minimal artificial selection for 10 generations in four locations, two organically and two conventionally managed sites, in Southern England.

## Material and Methods

### Creation of populations and description of locations

The CCPs were created by inter-crossing two sets of bread-wheat varieties: eight feed varieties and eleven milling varieties, plus the variety Bezostaya which was in both sets (Fig. <u>1</u> and Table S 1). F_1_ plants were self-fertilized and the number of F_2_ seeds from each cross counted and subsequently pooled. Seeds from 93 successful crosses (mean of 957 seeds per cross and range: 37 to 2569), entered the pooled founding population, subsequently termed FND. The pooled seeds were separated and sown by hand at each of the four locations in October 2003 in single plots (Fig. 1 and see also Döring et al., 2015). In subsequent generations, the populations were grown in a randomized complete block design (RCBD) with three replications with an average plot size of 25 m^2^ and a sowing density of 250 seeds/m^2^, giving an average demographic populations size of 32,000 plants. The seeds from each population were harvested and a proportion was re-sown each year at each location without any artificial selection.

**Fig. 1.**
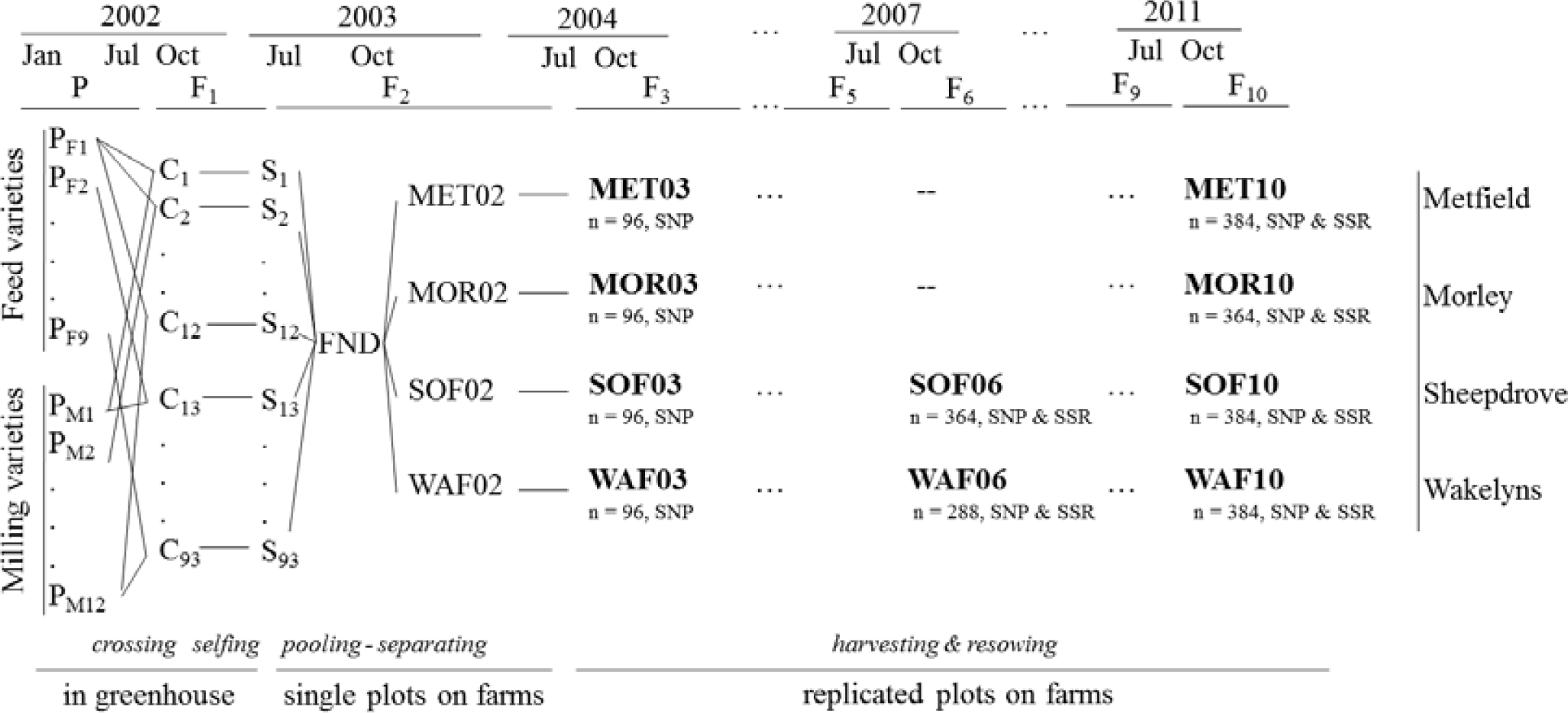
Schematic overview of the crossing scheme of the bread wheat CCP and of the sampled populations (bold fonts). The number of sampled individuals (n) and the sets of markers that were analyzed are shown below each population.

CCPs were grown at four locations: two organically managed: Wakelyns Agroforestry (WAF), in Suffolk (52° 39’N, 1° 17’E) and Sheepdrove Organic Farm (SOF) in Berkshire (51° 31’N, 1° 30’W); and two conventionally managed: Metfield Hall Farm (MET), directly adjacent to WAF in Suffolk (52° 41’N, 1° 29’E) and Morley Research Station (MOR) in Norfolk (52° 56’N, 1° 10’E). Fertilizer, pesticide and growth regulator applications at MET and MOR followed commercial practice. No pesticides were applied at WAF and SOF. At the organic locations weeds were controlled mechanically and through rotational design. For detailed descriptions of climatic and soil conditions see Jones et al. (2010) and Döring et al. (2015).

### Sampling of plant material and phenotyping

At SOF and WAF in generation 6 and 10, individual plants were tagged in the field, and plant height and heading date (day when ear is half-way emerged from flag leaf) were recorded. Individual plants were harvested, threshed, and grain yield of the whole plant was determined. From each plant, three random seeds were germinated and leaf tissue from one seedling was sampled for DNA extraction (final numbers of samples given in Fig. 1). At generation 3, from each location 150 seeds and at generation 6 and 10, 500 seeds were sampled from pooled plot harvests for genotyping. Sampling at generation 6 was only conducted at MET and MOR. DNA was extracted from leaf tissue (final sample number given in Fig. 1). As the genotype of the parental lines was crucial for the generation of the virtual FND (see below), DNA was extracted from five different seedlings per parental variety.

To evaluate the effect of the markers on pure stand performance, common plot trials with 19 of the total 20 parental varieties (except Norman) were assessed at the same four locations for plant height, obtained from 10 randomly chosen plants per plot, grain yield and yield components. These trials were conducted in three consecutive years (2005-2007) in a RCBD design with three replicates. For more detailed descriptions see Jones et al. (2010).

### Genotyping

After running a set of 70 publicly available SSR markers on the parental varieties and assessing the number of alleles and amplification quality, a subset of 18 SSR markers was chosen: 15 markers from Röder et al. (1998) (*gwm44, gwm46, gwm165, gwm186, gwm190, gwm213, gwm234, gwm325, gwm337, gwm469, gwm539, gwm583, gwm610 and gwm626*, of which *gwm44* and *gwm165* produced two loci), two markers from Stephenson et al. (1998) (*psp3100* and *psp3103*) and each one marker from Edwards et al. (1996) (*wmc56*) and Somers et al. (2004) (*barc134*). As two markers produced two loci, all-in-all 20 SSR loci were used for the genotype analysis.

Additionally, eight SNP markers were included, which were shown to be diagnostic for major genes involved in plant height (*Rht-B1* and *Rht-D1*), vernalization requirement (*Vrn-A1*), photoperiod response (*Ppd-A1, Ppd-B1, Ppd-D1,* and *Ppd-D1*(*D2*)) and one marker linked to the *1B/1R* chromosome translocation from rye (Zeller et al., 1973). Information on the SNP markers can be found at http://www.cerealsdb.uk.net/cerealgenomics/CerealsDB/kasp_download.php.

### Statistical analysis

#### Creation of a virtual founding population (FND)

As no seeds were kept from the original pooled founding population, the genotypic composition of this population could not be determined directly. Instead, we generated a virtual founding population by creating the heterozygous genotypes of each cross based on the genotype of the parentals. Subsequently we added the genotypes of each cross to the FND containing 10,000 individuals, proportionally to the recorded number of seeds that went into the ‘real’ founding population (see above). The final number of the genotype from the cross of line i and line j in the FND is thus 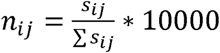, where, *s*_*ij*_ is the number of seeds from the cross of line i and line j, Σ *s*_*ij*_ is the sum of seeds from all crosses, and *n*_*ij*_ was rounded to the nearest integer. We compared this approach to mixing parental genotypes proportionally 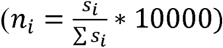, and to using the weight instead of the number of seeds (*s*_*ij*_as the weight of seeds) of each cross that went into the real founding population. The former approach of mixing parental genotypes proportionally showed no difference in allele frequencies. The approach based on seed weights had a very small impact on the resulting allele frequencies with a mean absolute difference of 0.005. For this reason, we only show the results for allele frequencies in which the FND was calculated based on seed number.

#### Treatment of SSR markers

At the SSR loci, alleles that were absent in parental genotypes (due to mutation or migration) were removed from the dataset, as the focus was on the assessment of allele frequencies. Mutations and migrations were considered as random and are thus assumed to have no biased effect on allele frequencies. To allow comparisons between the SSR and SNP marker sets and to avoid further assumptions in the mathematical treatment of multi-allelic markers (Goldringer and Bataillon, 2004; Meirmans and Hedrick, 2010), the marker data of the SSR markers were changed to bi-allelic markers. For each locus, the most frequent allele in the founding population was set as the first and all other alleles were combined into the second allele. The number of alleles and parental genotypes carrying the most frequent allele are shown in Table S 1.

#### Gene diversity

As a measure of genetic diversity within populations, we estimated Nei’s gene diversity (*H*_*e*_), which equals the expected heterozygosity under Hardy-Weinberg equilibrium (Nei, 1973). We calculated *H*_*e*_ for each locus and subsequently averaged over loci. 95% confidence intervals (CIs) were generated by bootstrapping over loci with 5,000 bootstraps, thereby avoiding specific assumptions about the distribution of the estimated parameters. Gene diversity was calculated with the R-package *hierfstat* (Goudet, 2005).

#### Effective population size and genetic differentiation

When testing the significance of changes in allele frequency due to selection it is important to test against changes due to genetic drift. Genetic drift is defined as the random change of allele frequencies resulting from the sampling of gametes over generations in a finite population (Hedrick, 2005). It can result in changes of allele frequency without any natural selection. The amount of genetic drift depends on the effective population size (*N*_*e*_), which is the number of fully outcrossing individuals in an ideal Wright-Fisher population undergoing the same rate of genetic change as the population under study (Wright, 1969). Note that *N*_*e*_ refers here to outcrossing individuals and does thus not refer directly to number of wheat plants. *N*_*e*_ can be derived by using neutral loci, which are assumed not to be subject to natural selection but only be affected by genetic drift. Subsequently, the expected amount of genetic drift can be calculated from the derived *N*_*e*_. Loci that have undergone positive or diversifying selection should then show increased, or respectively decreased, levels of change in allele frequency (Goldringer and Bataillon, 2004).

Genetic differentiation can also result from pure genetic drift. We therefore used the genetic differentiation of neutral loci at generation 10 to estimate *N*_*e*_. To remove loci under balancing or differential selection we employed the relation of expected genetic differentiation to heterozygosity (Beaumont and Nichols, 1996; Excoffier et al., 2009a). In particular, we used the function to detect loci under selection in the *Arlequin* package (Excoffier et al., 2009b), which simulates the expected *F*_*ST*_ null distribution under genetic drift in a finite island model, dependent on the expected heterozygosity. We ran 20,000 simulations with 100 demes. The remaining loci are thus assumed to be neutral regarding differential selection. For the calculation of *N*_*e*_ from observed *F*_*ST*_, we modified the equation from Hedrick (2005 p. 502), as *F*_*ST*_ is sampled from pairs of populations (see Supporting Method M2). Genetic differentiation between subpopulations with each size *N*_*e*_ after *t* generations of genetic drift without any migration between subpopulation is thus

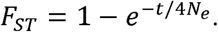

To estimate *N*_*e*_ from observed *F*_*ST*_, the equation needs to be solved for *N*_*e*_:

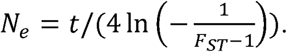

F_ST_ was estimated by Weir and Cockerham’s *F*_*ST*_ (Weir and Cockerham, 1984) as implemented in *hierfstat*, which is based on an analysis of molecular variance (AMOVA) comparing between population to within population diversity. As an error estimate for N_e_ we produced 95% confidence intervals (CI) for *F*_*ST*_ with the *boot.vc* function in the *hierfstat* package as suggested by Weir and Cockerham (1984), and subsequently calculated the CIs for N_e_. As we used the F_ST_ estimate from generation 10, and the populations were separated to the different locations in generation 2 (see Fig. 1), we used *t* = 8 as the number of generations since the populations could differentiate through genetic drift.

In order to compare the observed changes in allele frequency to the expected changes under pure genetic drift we calculated the 95% CIs of the allele frequency for each locus after *t* generations of genetic drift (Waples, 1989) as

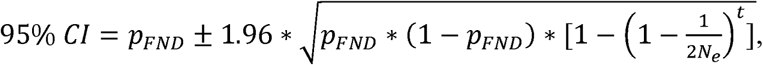

where *p*_*FND*_ is the allele frequency of the frequent allele in FND and *N*_*e*_ the effective populationsize. We furthermore compared this method to the temporal method as first proposed by Waples (1989), which is based on allele frequencies at two different generations (see Supporting Method M1).

#### Phenotypic effects

To investigate the phenotypic effects of the selected alleles, we carried out an association analysis with two sets of data: (1) phenotypic assessments on single plants in mixed stands, i.e. within the diverse populations described so far, and (2) phenotypic assessments in pure stands of single genotypes. The latter reflects common crop stands of single genotypes and was assessed in standard plot field trials. Whereas in mixed stands every plant is surrounded by different genotypes, in pure stands the whole plot or field consists of one single genotype.

The effects of the marker loci in single plants in mixed stands were assessed on the tagged plants (see above) at SOF and WAF in generation 6 and 10, i.e. in four field trials in total. Data were analyzed with a mixed model for each marker locus separately:

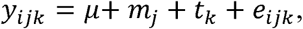

where *y*_*ijk*_ is the response of the i-th plant, carrying the j-th allele, in the k-th trial, *μ* the grand mean, *m*_*j*_ the effect of the j-th allele, *t*_*k*_ the effect of the k-th trial, and *e*_*ijk*_ are the residuals. The effect of the trial and the residuals were considered as random. Only individuals being homozygous for the respective marker were included.

The effects of the marker loci in pure stands, i.e. from the CCPs’ parent varieties, were assessed in common plot trials (3 years × 4 locations yielding overall 12 trials). First, adjusted means were calculated with the mixed model

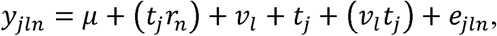

where *y*_*jln*_ is the response of the l-th variety in the n-th replicate block in the j-th trial, *μ* the grand mean,(*t*_*j*_*r*_*n*_) is the effect of the n-th replicate block in the j-th trial, *v*_*l*_ is the effect of l-th variety,*t*_*j*_ is the effect of the i-th trial, (*v*_*l*_*t*_*j*_) the variety-trial interaction and *e*_*jln*_ are the residuals. All effects except the variety effect were taken as random. Subsequently, the marker effect was assessed on the adjusted means with the following fixed model, where again one model was run for each marker locus:

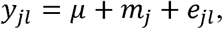

where *y*_*lj*_ is the response of the l-th variety carrying the j-th allele, *m*_*j*_ the effect of the j-th allele and *e*_*jl*_ are the residuals.

Significance of the marker effects were tested by an F-test against the residuals. The direction and size of the additive allele effect of the frequent allele was calculated as half the difference of the adjusted means of individuals homozygous for the frequent allele minus the adjusted means of individuals homozygous for the rare allele.

To relate the additive allele effect to the direction of selection we performed Pearson correlation between the additive allele effect and the difference in allele frequency between FND and generation 10. Significance of the correlation was tested with a t-test.

The statistical analysis was performed in R, version 3.5.0 (R Core Team, 2018). Bootstrapping was performed with the *boot* package (Canty and Ripley, 2016), linear models were fit with the *lme4* package (Bates et al., 2014), adjusted means were extracted with the *emmeans* package (Lenth, 2018), and ANOVA of the fixed effects was conducted with the *lmerTest* package (Kuznetsova et al., 2017).

## Results

### Gene diversity

All alleles that were present in the FND were also found in all sampled populations, so none of the alleles were eliminated over 10 generations. The two marker types show different levels of gene diversity (Fig. 2). However, as this measure depends on the allele frequencies the marker types cannot be directly compared. Diversity in the SSR set remained constant at 0.44, it decreased from 0.28 to 0.20 in the SNP marker set (Fig. 2). Estimates within generations did not differ between locations, indicating that the populations underwent similar changes at the four different locations.

**Fig. 2:**
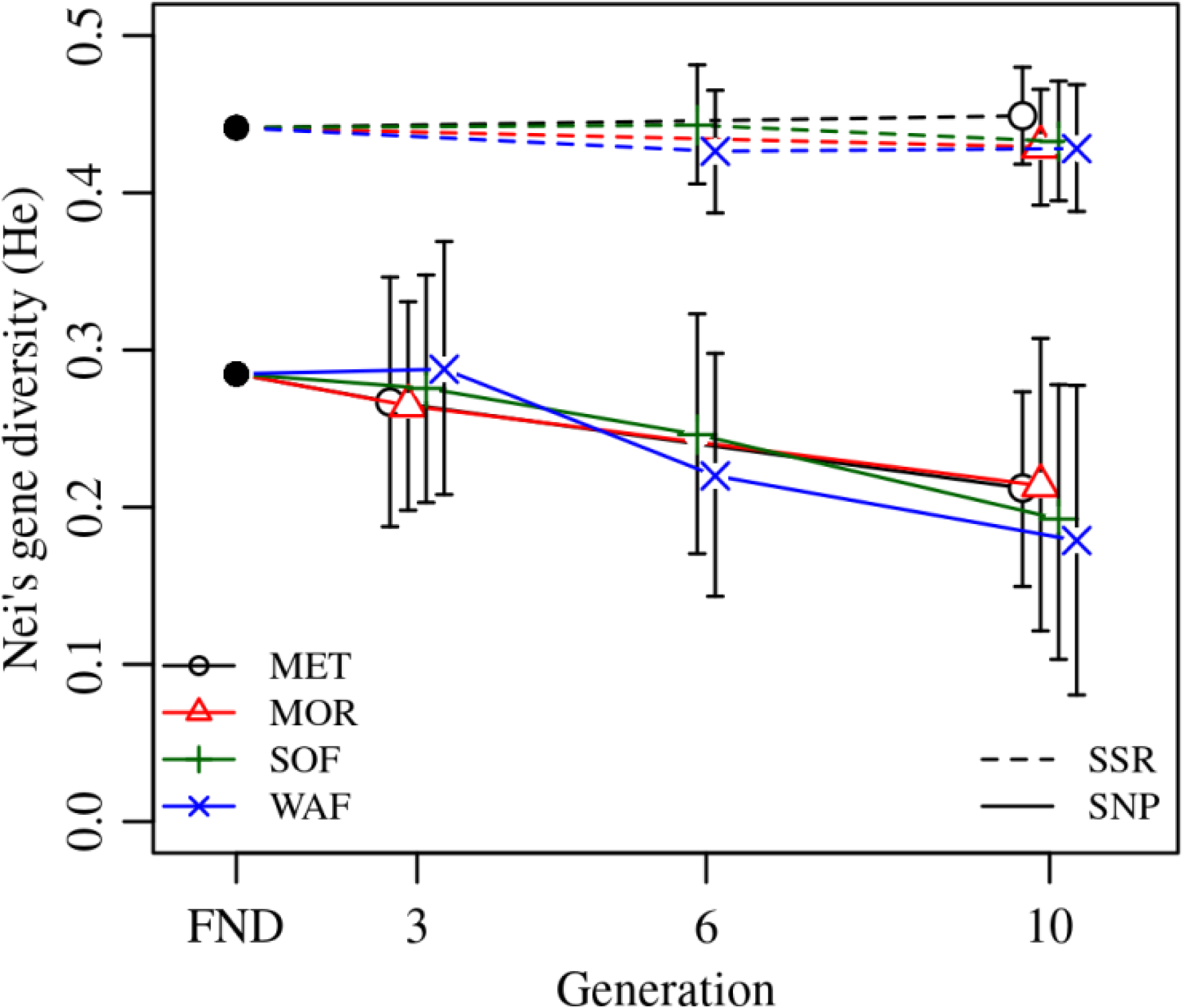
Change of Nei’s gene diversity (*H*_*e*_) over generations of bread wheat CCP for SNP and SSR marker sets at the four different locations Metfield (MET), Morley (MOR), Sheepdrove Organic Farm (SOF) and Wakelyns Agroforestry (WAF). FND indicates the founding population. Error bars are 95% CIs from bootstrapping over loci.

### Genetic differentiation

To investigate if the populations underwent a differential selection at the four locations, we estimated Weir and Cockerham’s *F*_*ST*_. *F*_*ST*_ of zero indicates no differentiation and one represents fixation. The overall differentiation at generation 10 using all marker loci *F*_*ST*_ was 0.013 [0.008-0.018]. Pairwise estimates (Table 1) range between *F*_*ST*_ = 0.006 and *F*_*ST*_ = 0.018. Both organically managed locations show a lower estimate of *F*_*ST*_ = 0.006 and a similarly small differentiation is found between SOF and MOR. To further test the differentiation due to management, we compared differentiation between both organically managed locations against both conventionally managed locations and compared this estimate to the pairwise groupings. The differentiation between management systems was higher (0.010 [0.005-0.015]) than both other groupings (WAF & MET vs. SOF & MOR: 0.005 [0.002-0.011] and SOF & MET vs. WAF & MOR: 0.005 [0.002-0.011]).

**Table 1.** Pairwise genetic differentiation at generation 10, measured by Weir and Cockerham’s *F*_*ST*_ (above diagonal) with 95% CIs from bootstrapping over loci (below diagonal). Metfield (MET) and Morley (MOR) are conventionally managed; Sheepdrove Organic Farm (SOF) and Wakelyns Agroforestry (WAF) are organically managed.

|  |  | conventional |  | organic |  |
| --- | --- | --- | --- | --- | --- |
|  |  | MET | MOR | SOF | WAF |
| conventional | MET |  | 0.011 | 0.015 | 0.018 |
|  | MOR | 0.005-0.020 |  | 0.006 | 0.012 |
| organic | SOF | 0.008-0.023 | 0.003-0.011 |  | 0.006 |
|  | WAF | 0.007-0.031 | 0.006-0.020 | 0.001-0.012 |  |

### Effective Population Size

Effective population size (*N*_e_) was estimated using the overall genetic differentiation in generation 10 and marker loci not under differential selection. These excluded three loci (*Ppd-B1, gwm165-4B*, and *gwm46-7B*), which were identified to be under differential selection (P<0.05 and *F*_*ST*_ higher than average, also see Figure S 1). Using the remaining loci, the overall genetic differentiation was estimated as *F*_*ST*_ = 0.009 [95% CI: 0.006-0.013], resulting in *N*_*e*_ = 221 [153-332]. The temporal method produced an estimate of *N*_*e*_ = 140, averaged across all comparisons for the SSR maker loci (Table S 2). Here, we report only the estimate from the SSR markers, as the investigation on gene diversity already indicates that selection took place in the SNP marker set, and absence of selection is a prerequisite for the temporal method.

### Changes in Allele Frequency

Given two estimates for *N*_*e*_, we inspected visually if the observed changes in allele frequency were greater than expected under pure genetic drift for *N*_*e*_ = 150 and *N*_*e*_ = 250 (Fig. 3). Instead of using only one final estimate, we used these two boundaries which allows comparisons of the size of expected genetic drift under different *N*_*e*_.

**Fig. 3:**
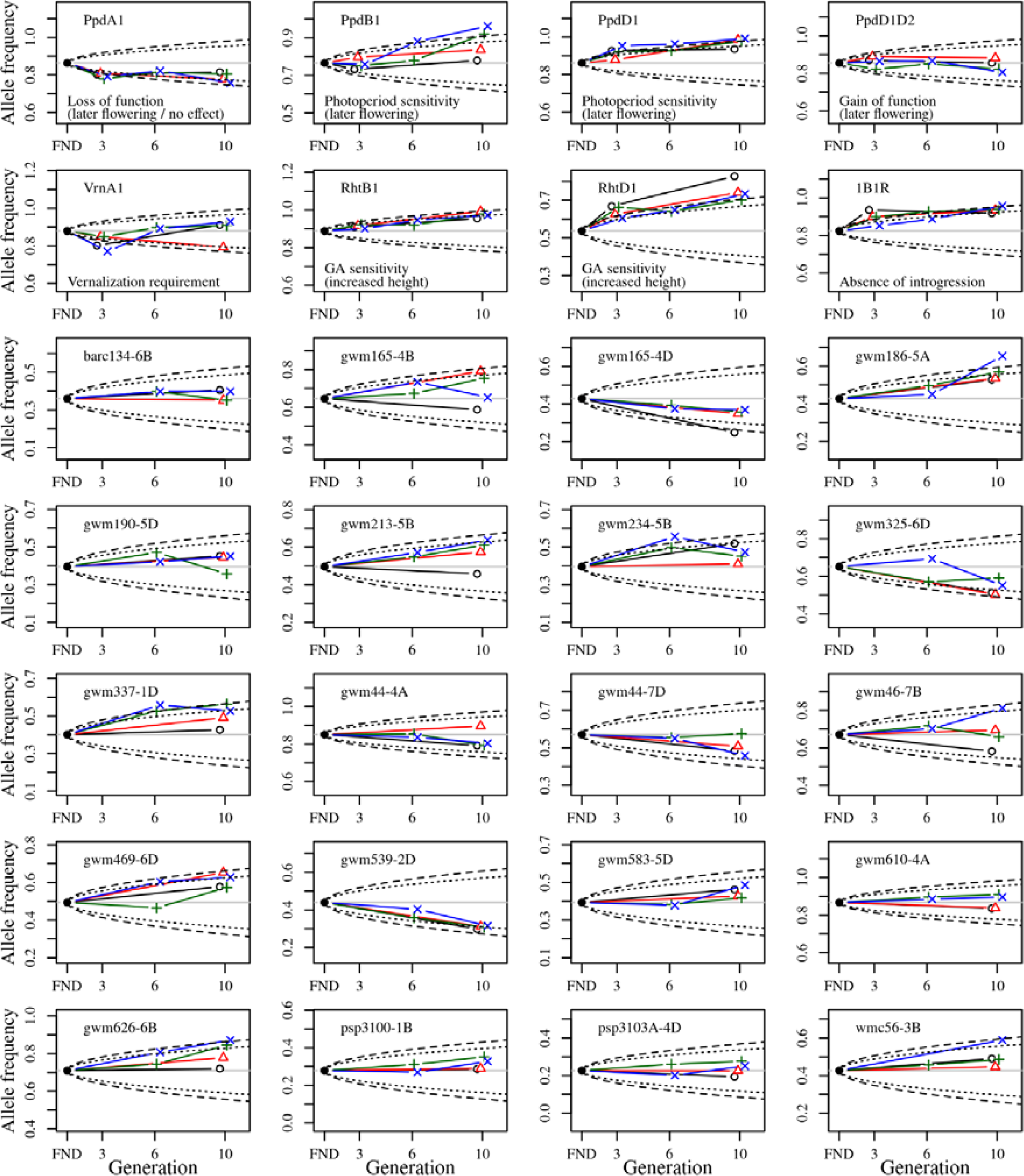
Change of allele frequency in the bread wheat CCP starting from the estimated allele frequency of the virtual founding population (FND). The allele frequency is shown for the frequent allele in the FND population. The different colors denote the allele frequencies in the populations at the different locations (black: MET, red: MOR, green: SOF, blue: WAF). The dashed and dotted lines indicate the 95% CI of the allele frequency expected under pure genetic drift given an *N*_*e*_ = 150 and *N*_*e*_ =250, respectively. For the SNP marker loci (top two rows), the function of the frequent allele is given.

Overall, the observed changes of allele frequency could occur due to genetic drift alone, even for the smaller *N*_*e*_ = 150. However, six of the gene-based markers (*Ppd-A1, Ppd-B1, Ppd-D1, Rht-B1, Rht-D1*, and *1B.1R*) show consistent changes over generations which are greater than expected under genetic drift. Except for *Ppd-B1*, the direction of selection was the same across locations and towards the wild type (WT) allele.

For *Ppd-B1* selection was only found at SOF and WAF. Furthermore, many SSR marker loci also showed similar selection for the same allele at all four locations (most notably *gwm165-4D, gwm186-5A, gwm539-2D*). The two loci which were identified to be under differential selection, *gwm165-4B*, and *gwm46-7B*, showed the greatest variation at generation 10, with selection at MET being different to the other locations. To test if selection was generally towards the similar direction at all four locations, we correlated the changes in allele frequency between FND and generation 10 at the different locations. Table 2 shows that the changes were highly correlated (P<0.001) between the different locations. The strongest correlation (r = 0.82) was between SOF and WAF, the two organically managed locations.

**Table 2:** Pearson correlation coefficients (above diagonal) of the difference of allele frequency between generation 10 at the different locations and the founding population (FND), with significance level indicated below the diagonal (***: P<0.001, based on a t-test with df = 26) for the four different locations Metfield (MET), Morley (MOR), Sheepdrove Organic Farm (SOF) and Wakelyns Agroforestry (WAF).

|  | MET | MOR | SOF | WAF |
| --- | --- | --- | --- | --- |
| MET |  | 0.70 | 0.64 | 0.70 |
| MOR | *** |  | 0.77 | 0.74 |
| SOF | *** | *** |  | 0.82 |
| WAF | *** | *** | *** |  |

### Phenotypic effects of the selected alleles

To investigate the phenotypic effects of the selected alleles, we correlated the changes of allele frequency with the additive allele effect in the mixed crop stands. Only plant height showed a significant correlation (P<0.01) to the overall changes of allele frequency between FND and generation 10 (Table 3 and Fig. 4). This relation indicates that height increasing alleles were under positive selection. This was true at both *Rht* homoeoloci. Interestingly, also *Ppd-A1* and *Ppd-D1* showed a significant effect on plant height (see F-Test for marker trait associations in Table S 3) again with the height increasing allele under positive selection (Fig. 4). Furthermore, at three SSR marker loci (*gwm165-4D, gwm325-6D*, and *gwm539-2D*), where there was consistent selection for the rare allele at all locations, the rare allele had a significant and increasing effect on plant height.

**Table 3:** Pearson correlations between the additive allele effects for the named traits measured in single plants in mixed stands and the change in allele frequency from FND to the average allele frequency at generation 10 (overall), and to the allele frequency at each location; *, ** denote significant correlation at P<0.05, and P<0.01, respectively.

| Allele effect on trait | Overall | MET | MOR | SOF | WAF |
| --- | --- | --- | --- | --- | --- |
| **Plant height** | 0.56** | 0.47* | 0.68*** | 0.51** | 0.36 |
| **Tillers per plant** | 0.30 | 0.02 | 0.23 | 0.46* | 0.38* |
| **Grain number per tiller** | -0.10 | -0.09 | -0.04 | -0.07 | -0.14 |
| **Thousand grain weight** | 0.23 | -0.02 | 0.24 | 0.42* | 0.20 |
| **Grain weight per tiller** | 0.18 | <0.01 | 0.22 | 0.35 | 0.11 |
| **Harvest index** | -0.35 | -0.22 | -0.36 | -0.40* | -0.29 |
| **Heading date** | 0.27 | 0.09 | 0.25 | 0.36 | 0.27 |

**Fig. 4:**
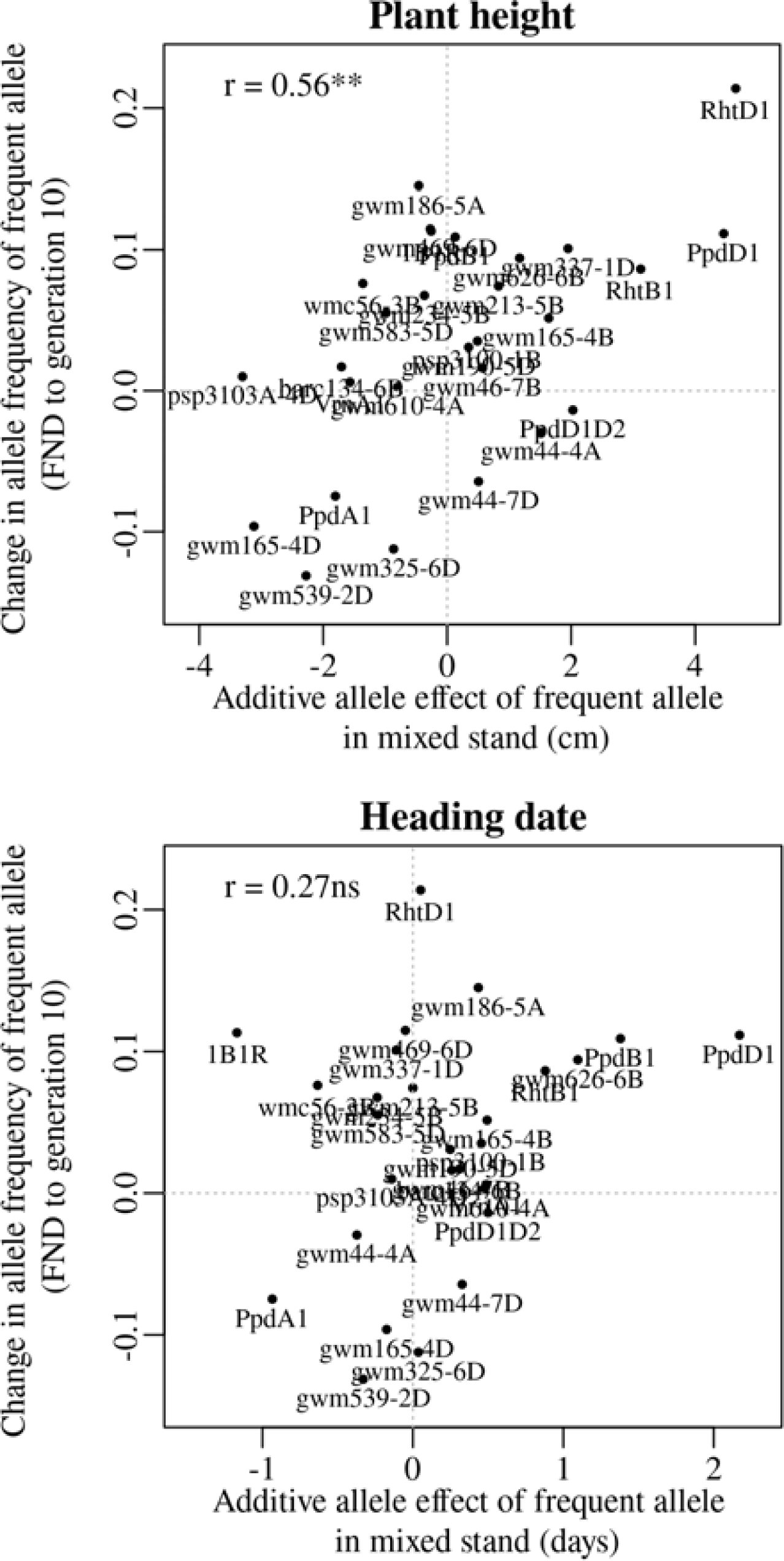
Relationship between the additive allele effect on plant height (left) and (b) on heading date (right) and the temporal change in allele frequency from the founding bread wheat CCP population (FND) to the allele frequency at generation 10 (averaged over all four locations). For plant height, the significant correlation indicates, that those genes with a stronger effect on plant height (such as *Rht-D1*) tended to have a more pronounced selection over time, demonstrated by the high change in allele frequency.

The relation between the additive allele effect on plant height and the change of allele frequency was also significant (P<0.01) at each location except at WAF (Table 3). At both organic locations, SOF and WAF, the change in allele frequency was significantly correlated with an increasing additive effect of the selected alleles on the number of tillers per plant. However, it is difficult to identify single markers, which are responsible for this significant correlation (Fig. 4**Error! Reference source not found.**). Interestingly, at the locus *gwm610-4A*, which shows the strongest effect on tillers per plant in single plants (Table S 3.), it has been selected – although not strongly – for the allele with increasing additive allele effect on tillers per plant at both organic locations and against this allele at both conventional locations (Fig. 3). Relationships between the additive allele effects and yield components (grain number per tiller, thousand grain weight and grain weight per tiller) were not significant neither were harvest index, and heading date were also not significant (Table 3).

Alleles often have pleiotropic effects, which can also result in the correlation between traits. As an example we calculated the correlations between plant height and various agronomically important traits. We compared these correlations among different traits for single plants in mixed stands compared to pure stands (Table 4). Increased plant height was associated with decreased grain number per tiller in pure stands, while this effect was not significant in the mixed stand. Similarly, plant height was negatively associated with grain weight per tiller in pure stands, while in mixed stands the relation was reversed, though not significant. For all three yield components (tillers per plant, grain number per tiller and thousand grain weight), the correlation with plant height was smaller (i.e. more negative or less positive) for the pure stands than for the mixed stands. In the mixed stands, where different genotypes directly competed, plant height was needed more to generate yield than in the pure stands.

**Table 4.**
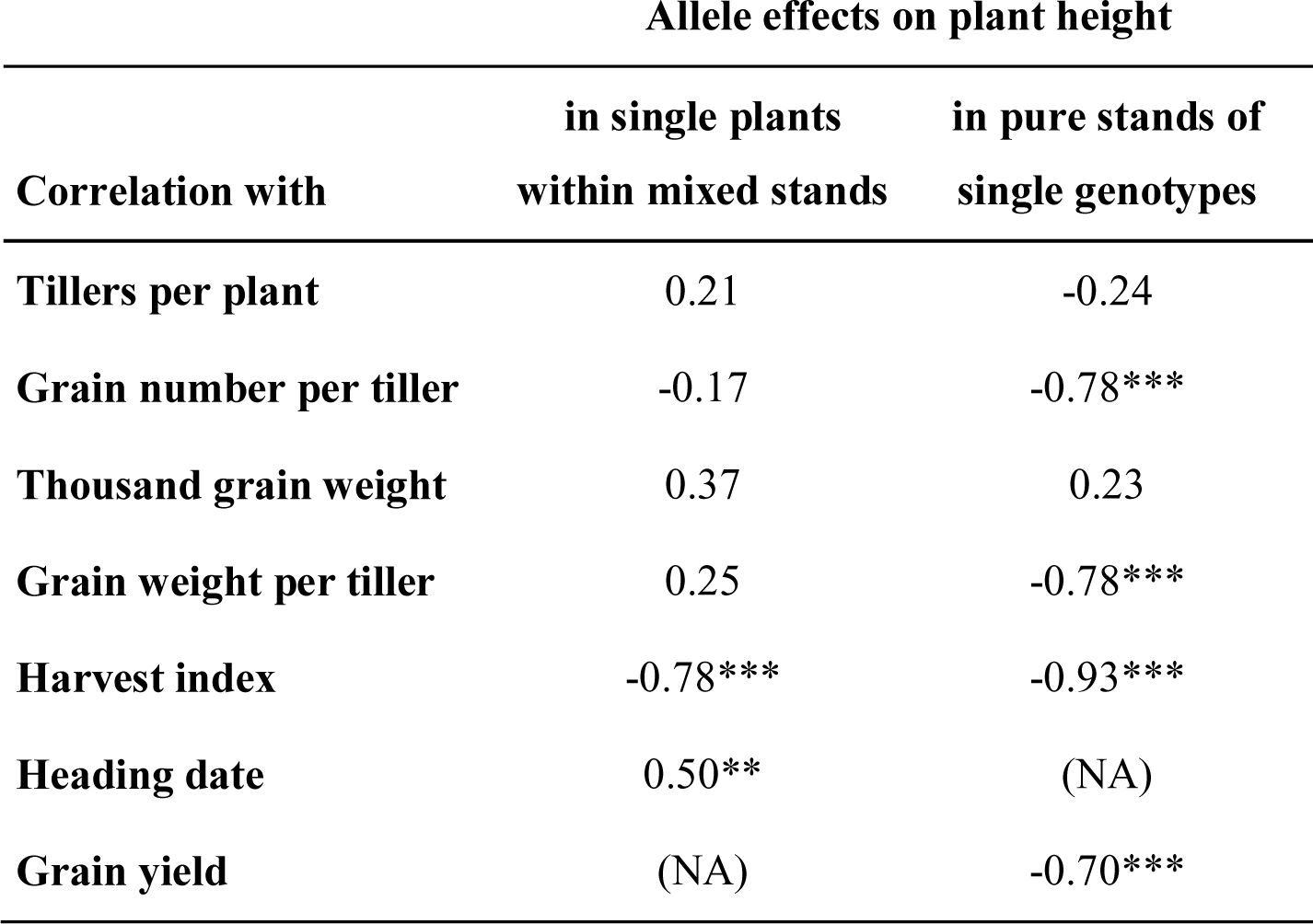
Correlations between the allele effects on plant height and allele effects on various yield components and heading date, for single plants within mixed stand and for pure stands of single genotypes; (NA): data not available; *, ** and *** denote significant correlations at P<0.05, P<0.01 and P<0.001, respectively.

| Correlation with | Allele effects on plant height |  |
| --- | --- | --- |
|  | in single plants<br>within mixed stands | in pure stands of<br>single genotypes |
| **Tillers per plant** | 0.21 | -0.24 |
| **Grain number per tiller** | -0.17 | -0.78*** |
| **Thousand grain weight** | 0.37 | 0.23 |
| **Grain weight per tiller** | 0.25 | -0.78*** |
| **Harvest index** | -0.78*** | -0.93*** |
| **Heading date** | 0.50** | (NA) |
| **Grain yield** | (NA) | -0.70*** |

Further analysis showed that grain yield (the product of the three yield components) in pure stand was mostly dependent on grain number per tiller (data not shown). However, natural selection acted towards a decreased grain number per tiller (Table 3).

## Discussion

### Gene diversity

While gene diversity did not decrease at the SSR markers in all four independent populations, gene diversity decreased at the SNP markers, with equal magnitude in all populations (Fig. 2). The fact that gene diversity remained constant at the SSR markers indicates that little or no selection had taken place on loci tracked by these markers. In contrast, the decrease in gene diversity at the SNP markers suggests that selection on these functional markers did take place, and that selection, overall, was of similar magnitude and direction at all locations. As absolute values of *H*_*e*_ were dependent on the genetic composition, they cannot be directly compared to other studies where populations of different composition were used. However, Raggi et al. (2016) using 22 SSR markers also found no decrease in *H*_*e*_ in a CCP of barley that had evolved for 13 years. More generally, our results confirm that overall genetic diversity in evolving wheat populations appears to be maintained to a large degree unless there is a strong specific selection force (e.g. Paillaird et al. (2000)).

### Effective population size

Estimation of effective population size (*N*_*e*_) is crucial for the identification of loci changes in allele frequency greater than expected under pure drift. The method based on the genetic differentiation (F_ST_) produced a higher estimate than the temporal method, using only the SSR markers (220 vs 140, Table S 2). The estimates from the temporal method based on the SNP markers showed an even smaller estimate (50, averaged over all comparisons), which is most likely due to selection taking place on these loci. As selection also took place on some of the SSR loci, the estimate from the temporal method also appears biased towards low estimates.

The values estimated in our study are higher than those reported by Rhoné et al.(2010) for different wheat populations grown over several generations in France, where estimates for *N*_e_ were 33, 114 and 118 at three locations. Other estimates from wheat populations in France, with *N*_e_ = 311 (Thépot et al., 2015) and *N*_e_ = 42 to *N*_e_ = 208 (Enjalbert et al., 1999) were closer to the values estimated here.

### Differentiation between locations

One of our main aims was to evaluate whether genetic differentiation occurred over ten generations for wheat CCPs growing conditions in contrasting environments with very different management. To assess the strength of population differentiation we took values between 0 and 0.05 as a general convention for little differentiation (Wright, 1978; Hartl and Clark., 1997). According to this convention, the values observed in this study are very low, sometimes not even significantly different from zero. Even when investigating *F*_*ST*_ values for the single loci, *F*_*ST*_ values were still below 0.05 (data not shown).

Although the locations in our study differed in management (organic vs conventional), resulting in strong variation in average yield level (around 9.5 t/ha at the conventional locations, and 5.3 t/ha at the organic locations (Jones et al., 2010; Döring et al., 2015), no consistent genetic differentiation regarding the management practices could be detected. In fact, pairwise comparisons between sites showed low values for genetic differentiation (Table 1), and selection was similar between locations, as indicated by the highly significant correlations between changes of allele frequencies (Table 2). In addition, markers diagnostic for genes of known function such as plant height and photoperiod sensitivity were not affected differently at the four studied locations (Fig. 3).

Three possible reasons for this lack of consistent genetic differentiation between the four locations are:

the number of generations was not sufficient to allow selection to exert a measurable effect on the genetic composition of the CCPs;

Some loci did exhibit environment specific selection but markers to detect these changes were not included in this study; Even though management is different between the locations, effective environmental conditions might be quite similar.

### Differentiation over time across all locations

The more exciting outcome of our analysis was the detection of a clear selection signature for in terms of temporal differentiation. At all four locations, genetic changes were observed in the same direction, in particular for the alleles linked to increased plant height and later flowering time (Table 3). These changes over time without any population differentiation can for most cases be summarized as selection towards wild-type. At the 5 loci for which significant changes of allele frequencies could be detected, there was selection for the wild-type alleles and against the mutant alleles that were introduced during the twentieth century through the implementation of sytstematic wheat breeding.

The two loci that have undergone the greatest change of allele frequencies were genes controlling height (*Rht-B1* and *Rht-D1*) (Fig. 3 and Fig. 4), suggesting that plant height has been a driving force in the evolutionary process of the investigated CCPs. This observation is supported by a study by Raquin et al. (2008), who found an increase of *Rht-B1* allele frequency in an experimental population of winter wheat from 0.66 in the initial generation to near-complete extinction of the dwarfing allele after 17 generations. The semi-dwarfing alleles at both loci were of major importance during the Green Revolution (Borlaug, 1983). Currently, 58 % of all European winter wheat varieties contain the *Rht-D1b* allele and 7% contain the *Rht-B1b* allele (Zanke et al., 2014). The selection for increased height can independently be observed in several other markers as well (Fig. 4, Table 3). These genetic effects confirm and explain the phenotypic observations on the same populations, which, in an earlier study, led to the conclusion that already in the third year of development, the wheat CCPs were almost 10 cm taller than the mean of the parents (Döring et al., 2015).

The selection for increased height may mainly be explained by competition for light. At the population level, competition in a genetically diverse plant stand selects for taller intra-specific competitors. It is therefore expected that genotypes with increased height are selected over time, and this is confirmed by our study. However, competition for light may not be the only driver for selection against the so-called dwarfing genes. In particular, the dwarfing genes *Rht-B1b* and *Rht-D1b* confer effects of reduced early vigour through shorter coleoptiles, redeuced viguor and in young plants. More generally, the selection observed in this study can be characterized as going towards more vegetative growth and more competitive ability, which could reduce yield potential(Denison, 2012).

The observed selection for increased height suggests that adaptation took place towards growing in a mixed stand population rather than to environmental conditions. This is because the dwarf genotypes, when grown together with taller neighbors will produce a reduced number of progenies, and thus be selected against over time. Thus, while the performance of a single plant in a mixed stand is determined by its competitive effect over its neighbours (e.g. through plant height), this is not the case in the pure stand.

The selection for the wild-type alleles at the genes *Ppd-B1L5* and *Ppd-D1* restores photoperiod sensitivity, as the mutant allele at both genes cause insensitivity to photoperiod (day length neutrality) and early heading. Again, these genes and alleles were very important for the Green Revolution (Borlaug, 1983), allowing very wide adaptation. Thus, also for the two *Ppd-1* genes, selection seems to have happened towards wild type and against alleles that are important in modern agricultural production. It should be noted also that the majority of UK wheat varieties are photoperiod sensitive so these results are alsoa reversion to UK type. Interestingly, the mutant allele of *Ppd1* is also responsible for a further shortening of plant height, due to the temporal shortening of vegetative growth (Börner et al., 1993). Accordingly, we observed a significant correlation between allele effects on plant height and heading date (Table 4, Fig. 4). Because of the pleiotropic effects it is still open, what the driving factor for the changes in allele frequency of the *Ppd* genes were in the wheat CCPs. In particular, it remains open whether later flowering itself conveyed a fitness advantage to individuals within the evolving wheat populations.

The selection for the wild type form can also be hypothesized for the *X1B.1R* marker. This marker identifies the translocation from rye into wheat, which is widespread in many breeding programs, and mostly originates from the rye variety Petkus (Schneider and Molnár-Láng, 2009). In the studied populations, selection occurred against the translocation, even though it is assumed that the introgression confers increased disease resistance (Heslop-Harrison et al., 1990) and should thus also lead to improved fitness under disease pressure.

Our observation is that the major pattern of selection is the result of selection for wild-type alleles, it may be suggested that evolutionary plant breeding approaches can be improved by fixing these alleles, so that negative selection cannot occur. Consequently, individual plants within the diverse population would not invest resources on competitional behaviour and selection would then be directed towards prevailing local growing conditions. However, traits that are linked to the competitive ability of the plant, such as plant height, are governed by a large number of genes (Zanke et al., 2014). It is therefore unlikely that competition within a population can be fixed genetically without substantially reducing genetic diversity. Since it seems impossible to create populations that are completely free of any trade-offs, future research will need to address the question which trade-offs show the greatest opportunities for developing multifunctional, and potentially adaptive, CCPs.

## Acknowledgements

The work was funded under the LINK scheme with financial support from the UKs Department of Environment Food and Rural Affairs (Defra), the Biotechnology and Biological Sciences Research Council (BBSRC), and the Agriculture and Horticulture Development Board (AHDB). The project title was; Adaptive winter wheat populations: development, genetic characterization and application (LK0999).

## Author contributions

SK performed data analysis and led writing of manuscript, TFD and HEJ conducted field experiments, MSW and JWS planned the experiments and won funding, LUW supported data analysis, MLW carried out molecular work, SG led project.

## References

Allard, R.W. 1988. Genetic Changes Associated with the Evolution of Adaptedness in Cultivated Plants and Their Wild Progenitors. J Hered 79(4): 225–238.

Anten, N.P.R., and P.J. Vermeulen. 2016. Tragedies and Crops: Understanding Natural Selection To Improve Cropping Systems. Trends in Ecology & Evolution 31(6): 429–439. doi: 10.1016/j.tree.2016.02.010.

Atlin, G.N., J.E. Cairns, and B. Das. 2017. Rapid breeding and varietal replacement are critical to adaptation of cropping systems in the developing world to climate change. Global Food Security 12: 31–37. doi: 10.1016/j.gfs.2017.01.008.

Bates, D., M. Maechler, B. Bolker, and S. Walker. 2014. lme4: Linear mixed-effects models using Eigen and S4. R package version 1.1-7. http://CRAN.R-project.org/package=lme4.

Beaumont, M.A., and R.A. Nichols. 1996. Evaluating loci for use in the genetic analysis of population structure. Proc. R. Soc. Lond. B 263(1377): 1619–1626. doi: 10.1098/rspb.1996.0237.

Bertholdsson, N.O., O. Weedon, S. Brumlop, and M.R. Finckh. 2016. Evolutionary changes of weed competitive traits in winter wheat composite cross populations in organic and conventional farming systems. European Journal of Agronomy 79: 23–30. doi: 10.1016/j.eja.2016.05.004.

Borlaug, N.E. 1983. Contributions of Conventional Plant Breeding to Food Production. Science 219(4585): 689–693. doi: 10.1126/science.219.4585.689.

Börner, A., A.J. Worland, J. Plaschke, E. Schumann, and C.N. Law. 1993. Pleiotropic Effects of Genes for Reduced Height (Rht) and Day-Length Insensitivity (Ppd) on Yield and its Components for Wheat Grown in Middle Europe. Plant Breeding 111(3): 204–216. doi: 10.1111/j.1439-0523.1993.tb00631.x.

Brown, J.K.M., and M.S. Hovmøller. 2002. Aerial Dispersal of Pathogens on the Global and Continental Scales and Its Impact on Plant Disease. Science 297(5581): 537–541. doi: 10.1126/science.1072678.

Canty, A., and B. Ripley. 2016. boot: Bootstrap R (S-Plus) Functions. R package version 1.3-18.

Cooper, M., and G.L. Hammer. 1996. Synthesis of strategies for crop improvement. p. 591–623. In Cooper, M., Hammer, G.L. (eds.), Plant Adaptation and Crop Improvement. CAB International, Wallingford, UK.

Denison, R.F. 2012. Darwinian agriculture: How understanding evolution can improve agriculture. Princeton University Press.

Döring, T.F., P. Annicchiarico, S. Clarke, Z. Haigh, H.E. Jones, H. Pearce, J. Snape, J. Zhan, and M.S. Wolfe. 2015. Comparative analysis of performance and stability among composite cross populations, variety mixtures and pure lines of winter wheat in organic and conventional cropping systems. Field Crops Research 183: 235–245. doi: 10.1016/j.fcr.2015.08.009.

Döring, T.F., S. Knapp, G. Kovacs, K. Murphy, and M.S. Wolfe. 2011. Evolutionary Plant Breeding in Cereals - Into a New Era. Sustainability 3(10): 1944–1971. doi: 10.3390/su3101944.

Edwards, K., J. Barker, A. Daly, C. Jones, and A. Karp. 1996. Microsatellite libraries enriched for several microsatellite sequences in plants. Biotechniques 20(5): 758.

Enjalbert, J., I. Goldringer, S. Paillard, and P. Brabant. 1999. Molecular markers to study genetic drift and selection in wheat populations. J Exp Bot 50(332): 283–290. doi: 10.1093/jxb/50.332.283.

Excoffier, L., T. Hofer, and M. Foll. 2009a. Detecting loci under selection in a hierarchically structured population. Heredity 103(4): 285–298. doi: 10.1038/hdy.2009.74.

Excoffier, L., G. Laval, and S. Schneider. 2009b. Arlequin: An Intergrated Software Package for Population Genetics Data Analysis, Version 3.5. Institute of Zoology, University of Berne, Berne, Switzerland.

Finckh, M.R., and M.S. Wolfe. 2006. Diversification strategies. p. 269–307. *In* Cooke, B.M., Jones, D.G., Kaye, B. (eds.), The Epidemiology of Plant Diseases. Springer Netherlands.

Gabriel, D., S.M. Sait, W.E. Kunin, and T.G. Benton. 2013. Food production vs. biodiversity: comparing organic and conventional agriculture. Journal of Applied Ecology 50(2): 355–364. doi: 10.1111/1365-2664.12035.

Goldringer, I., and T. Bataillon. 2004. On the Distribution of Temporal Variations in Allele Frequency. Genetics 168(1): 563–568. doi: 10.1534/genetics.103.025908.

Goudet, J. 2005. hierfstat, a package for R to compute and test hierarchical F-statistics. Molecular Ecology Notes 5(1): 184–186. doi: 10.1111/j.1471-8286.2004.00828.x.

Hartl, D.L., and A.. Clark. 1997. Principles of Population Genetics. Sinauer Associates, Inc, Sunderland, MA.

Hedrick, P.W. 2005. Genetics of Populations. Jones and Bartlett Publishers, Sudbury, Massachusettes.

Heslop-Harrison, J.S., A.R. Leitch, T. Schwarzacher, and K. Anamthawat-Jonsson. 1990. Detection and characterization of 1B/1R translocations in hexaploid wheat. Heredity 65(3): 385–392.

Jones, H., S. Clarke, Z. Haigh, H. Pearce, and M. Wolfe. 2010. The Effect of the Year of Wheat Variety Release on Productivity and Stability of Performance on Two Organic and Two Non-Organic Farms. The Journal of Agricultural Science 148(03): 303–317. doi: 10.1017/S0021859610000146.

Kuznetsova, A., P.B. Brockhoff, and R.H.B. Christensen. 2017. lmerTest: Tests in Linear Mixed Effects Models. Journal of Statistical Software 82(13): 1–26. doi: 10.18637/jss.v082.i13.

Le Boulc’h, V., J.L. David, P. Brabant, and C. de Vallavieille-Pope. 1994. Dynamic conservation of variability: responses of wheat populations to different selective forces including powdery mildew. Genet Sel Evol 26(S1): S221. doi: 10.1186/1297-9686-26-S1-S221.

Lenth, R. 2018. emmeans: Estimated Marginal Means, aka Least-Squares Means.

Litrico, I., and C. Violle. 2015. Diversity in Plant Breeding: A New Conceptual Framework. Trends in Plant Science 20(10): 604–613. doi: 10.1016/j.tplants.2015.07.007.

Meirmans, P.G., and P.W. Hedrick. 2010. Assessing population structure: FST and related measures. Molecular Ecology Resources 11(1): 5–18. doi: 10.1111/j.1755-0998.2010.02927.x.

Mercer, K.L., and H.R. Perales. 2010. Evolutionary response of landraces to climate change in centers of crop diversity. Evolutionary Applications 3(5–6): 480–493. doi: 10.1111/j.1752-4571.2010.00137.x.

Nei, M. 1973. Analysis of Gene Diversity in Subdivided Populations. Proceedings of the National Academy of Sciences of the United States of America 70(12): 3321–3323.

Nei, M., and F. Tajima. 1981. Genetic Drift and Estimation of Effective Population Size. Genetics 98(3): 625–640.

Paillard, S., I. Goldringer, J. Enjalbert, G. Doussinault, C. de Vallavieille-Pope, and P. Brabant. 2000. Evolution of resistance against powdery mildew in winter wheat populations conducted under dynamic management. I-Is specific seedling resistance selected? TAG Theoretical and Applied Genetics 101(3): 449–456.

R Core Team. 2018. R: A Language and Environment for Statistical Computing. R Foundation for Statistical Computing, Vienna, Austria.

Raggi, L., S. Ceccarelli, and V. Negri. 2016. Evolution of a barley composite cross-derived population: an insight gained by molecular markers. The Journal of Agricultural Science 154(01): 23–39. doi: 10.1017/S0021859614001269.

Raquin, A.-L., P. Brabant, B. Rhoné, F. Balfourier, P. Leroy, and I. Goldringer. 2008. Soft selective sweep near a gene that increases plant height in wheat. Molecular Ecology 17(3): 741–756. doi: 10.1111/j.1365-294X.2007.03620.x.

Rhoné, B., R. Vitalis, I. Goldringer, and I. Bonnin. 2010. Evolution of flowering time in experimental wheat populations: a comprehensive approach to detect genetic signatures of natural selection. Evolution 64(7): 2110–2125. doi: 10.1111/j.1558-5646.2010.00970.x.

Röder, M.S., V. Korzun, K. Wendehake, J. Plaschke, M.-H. Tixier, P. Leroy, and M.W. Ganal. 1998. A Microsatellite Map of Wheat. Genetics 149(4): 2007–2023.

Schneider, A., and M. Molnár-Láng. 2009. Detection of the 1RS chromosome arm in Martonvásár wheat genotypes containing 1Bl.1Rs or 1Al.1Rs translocations using SSR and STS markers. Acta Agronomica Hungarica 57(4): 409–416. doi: 10.1556/AAgr.57.2009.4.3.

Somers, D.J., P. Isaac, and K. Edwards. 2004. A high-density microsatellite consensus map for bread wheat (Triticum aestivum L.). Theoretical and Applied Genetics 109: 1105–1114. doi: 10.1007/s00122-004-1740-7.

Stephenson, P., G. Bryan, J. Kirby, A. Collins, K. Devos, C. Busso, and M. Gale. 1998. Fifty new microsatellite loci for the wheat genetic map. TAG Theoretical and Applied Genetics 97: 946–949. doi: 10.1007/s001220050975.

Suneson, C.A. 1956. An Evolutionary Plant Breeding Method. Agron J 48(4): 188–191.

Thépot, S., G. Restoux, I. Goldringer, F. Hospital, D. Gouache, I. Mackay, and J. Enjalbert. 2015. Efficiently Tracking Selection in a Multiparental Population: The Case of Earliness in Wheat. Genetics 199(2): 609–623. doi: 10.1534/genetics.114.169995.

Waples, R.S. 1989. A Generalized Approach for Estimating Effective Population Size From Temporal Changes in Allele Frequency. Genetics 121(2): 379–391.

Weiner, J. 2003. Ecology–the science of agriculture in the 21st century. The Journal of Agricultural Science 141(3–4): 371–377.

Weiner, J., S.B. Andersen, W.K.-M. Wille, H.W. Griepentrog, and J.M. Olsen. 2010. Evolutionary Agroecology: the potential for cooperative, high density, weed-suppressing cereals. Evolutionary Applications 3(5–6): 473–479. doi: 10.1111/j.1752-4571.2010.00144.x.

Weiner, J., and R.P. Freckleton. 2010. Constant Final Yield. Annual Review of Ecology, Evolution, and Systematics 41(1): 173–192. doi: 10.1146/annurev-ecolsys-102209-144642.

Weir, B., and C. Cockerham. 1984. Estimating F-statistics for the analysis of population structure. Evolution 38(6): 1358–1370. doi: 10.2307/2408641.

Wright, S. 1969. Evolution and the Genetics of Populations: Volume 2, The Theory of Gene Frequencies. University of Chicago Press, Chicago.

Wright, S. 1978. Evolution and the Genetics of Population, Variability Within and Among Natural Populations. University of Chicago Press, Chicago.

Zanke, C.D., J. Ling, J. Plieske, S. Kollers, E. Ebmeyer, V. Korzun, O. Argillier, G. Stiewe, M. Hinze, K. Neumann, M.W. Ganal, and M.S. Röder. 2014. Whole Genome Association Mapping of Plant Height in Winter Wheat (Triticum aestivum L.) (P Hernandez, Ed.). PLoS ONE 9(11): e113287. doi: 10.1371/journal.pone.0113287.

Zeller, F.J., E.R. Sears, and L.M.S. Sears. 1973. 1B/1R wheat-rye chromosome substitutions and translocations. p. 209–221. *In* Proceedings of the fourth international wheat genetics symposium. Alien genetic material. University of Missouri.

