## Supplementary data for "Natural selection towards wild-type in composite cross populations of winter wheat"

**Supporting Information**

***Supporting Methods***

*M1: Estimation of Ne with the 'temporal method'*

As a second method to estimate *N*_e_, we used the 'temporal method' as first developed by Waples (1989), which is based on the allele frequencies in two generations. We first calculated the standardized variance of gene frequency change, $\hat{F_{c}}$, for each locus, assuming sampling before reproduction (eq. 8 in Waples (1989)):

$\hat{F_{c}}= \frac{1}{K}\sum_{i=1}^{K} \frac{{(x_{i}-y_{i})}^{2}}{(x_{i}-y_{i})/2-x_{i}y_{i}}$,

where K is the number of alleles, $x_{i}$and $y_{i}$ are the frequencies of the i-th allele in the two different generations. As all loci had two alleles (see Material and Methods in the main section) the genomewide-estimate ($\bar{F_{c}}$) was calculated as the mean over all per loci estimates. The number of sampled individuals was calculated as harmonic mean over loci (Nei and Tajima, 1981) and Ne was calculated as

$N_{e}=\frac{t}{2[\bar{F_{c}}-\frac{1}{2S_{1}}-\frac{1}{2S2}]}$*,*

where $t$ is the number of generations, and $S_{1}$and $S_{2}$are the numbers of sampled individuals in the first and in the second generation, respectively. 95% confidence intervals (CI) for the estimates of $N_{e}$ were derived, assuming that $\frac{n\bar{F_{c}}}{E(\bar{F_{c}})}$ is *χ^2^* –distributed with $n$ degrees of freedom and $n$ being the numbers of loci (Lewontin and Krakauer, 1973)

$$95\% CI \left( \bar{F_{c}} \right)=\{\frac{n\bar{F_{c}}}{\chi^{2}\left( 0.025,n \right)};\frac{n\bar{F_{c}}}{\chi^{2}\left( 0.975,n \right)}\}$$

and subsequent calculation the 95% CIs of $N_{e}$.

***Supporting Tables***

Table S 1. Chromosome, number of alleles scored in the parental genotypes and parental genotypes carrying the most frequent alleles. A one indicates that this parental genotype carries the most frequent allele for the given locus.

| **Locus** | **Marker set** | **Chromosome** | **Number of alleles** | **Bezostaya** | **Buchan** | **Cadenza** | **Claire** | **Deben** | **Hereward** | **HTL** | **Mercia** | **Monopol** | **MWidgeon** | **Norman** | **Option** | **Pastiche** | **Renan** | **Renesansa** | **Soissons** | **Spark** | **Tanker** | **Thatcher** | **Wembley** |
| --- | --- | --- | --- | --- | --- | --- | --- | --- | --- | --- | --- | --- | --- | --- | --- | --- | --- | --- | --- | --- | --- | --- | --- |
| B1R_1B | SNP | 1B | 2 | 1 | 0 | 1 | 1 | 1 | 1 | 0 | 1 | 1 | 1 | 1 | 1 | 1 | 1 | 1 | 1 | 1 | 0 | 1 | 1 |
| Ppd-A1Cdex_2A | SNP | 2A | 2 | 1 | 1 | 1 | 1 | 1 | 1 | 1 | 1 | 1 | 0 | 1 | 1 | 1 | 1 | 1 | 1 | 0 | 1 | 1 | 1 |
| Ppd-B1L5_2B | SNP | 2B | 2 | 0 | 1 | 1 | 1 | 1 | 1 | 1 | 1 | 1 | 1 | 1 | 1 | 1 | 0 | 1 | 0 | 1 | 1 | 0 | 0 |
| Ppd-D1_2D | SNP | 2D | 2 | 0 | 1 | 1 | 1 | 1 | 1 | 1 | 1 | 1 | 1 | 1 | 1 | 1 | 1 | 0 | 0 | 1 | 1 | 1 | 1 |
| Ppd-D1D2_2D | SNP | 2D | 2 | 1 | 1 | 1 | 1 | 0 | 1 | 1 | 1 | 1 | 1 | 0 | 1 | 0 | 1 | 1 | 1 | 1 | 1 | 1 | 1 |
| Rht-B1_4B | SNP | 4B | 2 | 1 | 1 | 1 | 1 | 1 | 1 | 1 | 1 | 1 | 1 | 1 | 1 | 1 | 0 | 1 | 0 | 1 | 1 | 1 | 1 |
| Rht-D1_4D | SNP | 4D | 2 | 1 | 0 | 1 | 0 | 0 | 0 | 1 | 1 | 1 | 1 | 0 | 0 | 0 | 1 | 1 | 1 | 1 | 0 | 1 | 0 |
| VrnA1prom_5A | SNP | 5A | 2 | 1 | 1 | 0 | 1 | 1 | 1 | 1 | 1 | 1 | 1 | 1 | 1 | 1 | 1 | 1 | 1 | 1 | 1 | 0 | 0 |
| barc134_6B | SSR | 6B | 7 | 0 | 1 | 0 | 0 | 1 | 1 | 0 | 0 | 0 | 0 | 1 | 1 | 0 | 0 | 0 | 0 | 0 | 1 | 0 | 1 |
| gwm165_4B | SSR | 4B | 4 | 0 | 1 | 1 | 0 | 1 | 1 | 1 | 1 | 0 | 1 | 1 | 0 | 1 | 0 | 0 | 0 | 1 | 1 | 1 | 1 |
| gwm165_4D | SSR | 4D | 4 | 1 | 1 | 0 | 0 | 1 | 0 | 0 | 1 | 0 | 0 | 1 | 1 | 1 | 0 | 1 | 0 | 0 | 0 | 0 | 1 |
| gwm186_5A | SSR | 5A | 5 | 0 | 1 | 0 | 1 | 1 | 1 | 0 | 0 | 1 | 1 | 0 | 1 | 0 | 0 | 0 | 1 | 0 | 0 | 0 | 0 |
| gwm190_5D | SSR | 5D | 4 | 1 | 0 | 1 | 0 | 0 | 1 | 0 | 0 | 0 | 1 | 0 | 0 | 0 | 1 | 1 | 1 | 1 | 0 | 0 | 0 |
| gwm213_5B | SSR | 5B | 6 | 0 | 1 | 0 | 1 | 1 | 1 | 1 | 1 | 0 | 1 | 1 | 0 | 1 | 0 | 0 | 0 | 0 | 0 | 1 | 0 |
| gwm234_5B | SSR | 5B | 5 | 0 | 1 | 0 | 1 | 1 | 0 | 1 | 1 | 0 | 0 | 0 | 1 | 1 | 0 | 0 | 0 | 0 | 0 | 0 | 0 |
| gwm325_6D | SSR | 6D | 4 | 1 | 1 | 0 | 1 | 1 | 1 | 1 | 1 | 1 | 0 | 1 | 0 | 1 | 1 | 0 | 0 | 1 | 0 | 0 | 1 |
| gwm337_1D | SSR | 1D | 5 | 0 | 0 | 1 | 0 | 1 | 0 | 1 | 1 | 0 | 1 | 1 | 0 | 1 | 1 | 0 | 0 | 0 | 0 | 0 | 0 |
| gwm44_4A | SSR | 4A | 2 | 1 | 1 | 1 | 0 | 0 | 1 | 1 | 1 | 1 | 1 | 1 | 1 | 1 | 1 | 1 | 1 | 1 | 1 | 1 | 1 |
| gwm44_7D | SSR | 7D | 4 | 0 | 1 | 1 | 0 | 0 | 1 | 0 | 0 | 1 | 1 | 1 | 0 | 1 | 1 | 0 | 1 | 1 | 1 | 0 | 1 |
| gwm46_7B | SSR | 7B | 4 | 0 | 1 | 1 | 1 | 1 | 0 | 1 | 1 | 0 | 1 | 1 | 0 | 0 | 0 | 1 | 0 | 1 | 1 | 1 | 0 |
| gwm469_6D | SSR | 6D | 5 | 0 | 1 | 1 | 0 | 1 | 1 | 1 | 0 | 1 | 0 | 0 | 1 | 0 | 0 | 0 | 0 | 1 | 0 | 1 | 1 |
| gwm539_2D | SSR | 2D | 5 | 0 | 0 | 0 | 1 | 1 | 0 | 0 | 1 | 0 | 0 | 1 | 0 | 0 | 0 | 0 | 1 | 0 | 1 | 1 | 1 |
| gwm583_5D | SSR | 5D | 4 | 1 | 0 | 0 | 0 | 0 | 1 | 1 | 1 | 1 | 0 | 0 | 1 | 0 | 1 | 0 | 0 | 1 | 0 | 0 | 0 |
| gwm610_4A | SSR | 4A | 2 | 1 | 1 | 1 | 1 | 1 | 1 | 1 | 1 | 1 | 1 | 1 | 1 | 1 | 1 | 1 | 0 | 1 | 1 | 0 | 0 |
| gwm626_6B | SSR | 6B | 2 | 0 | 1 | 1 | 1 | 1 | 1 | 1 | 1 | 1 | 1 | 1 | 1 | 0 | 0 | 0 | 0 | 1 | 1 | 0 | 0 |
| psp3100_1B | SSR | 1B | 7 | 0 | 0 | 0 | 1 | 1 | 0 | 0 | 0 | 0 | 1 | 1 | 0 | 1 | 0 | 0 | 0 | 0 | 0 | 0 | 0 |
| psp3103A_4D | SSR | 4D | 7 | 0 | 0 | 0 | 0 | 1 | 1 | 0 | 0 | 0 | 0 | 1 | 1 | 1 | 0 | 0 | 0 | 0 | 0 | 0 | 0 |
| wmc56_3B | SSR | 3B | 5 | 0 | 1 | 0 | 1 | 1 | 0 | 1 | 0 | 0 | 0 | 1 | 1 | 0 | 1 | 0 | 0 | 0 | 0 | 0 | 0 |

Table S 2: Estimates of effective population size based on the temporal method as described in supporting methods (M1).

| **Marker set** | **Paired generations** | **FND vs  generation 6** | | **FND vs  generation 10** | | | | **Generation 6  vs 10** | |
| --- | --- | --- | --- | --- | --- | --- | --- | --- | --- |
|  | **Site** | **SOF** | **WAF** | **MET** | **MOR** | **SOF** | **WAF** | **SOF** | **WAF** |
| SSR | Ne | 210 | 128 | 156 | 165 | 146 | 88 | 144 | 74 |
|  | 95% CI | [94-393] | [58-237] | [73-280] | [77-293] | [68-259] | [41-154] | [62-288] | [33-139] |
| SNP | Ne | 78 | 47 | 56 | 40 | 43 | 29 | 57 | 56 |
|  | 95% CI | [25-226] | [15-135] | [17-141] | [12-99] | [13-109] | [9-73] | [15-137] | [14-137] |

Table S 3. F-Test of the marker main effect in single plants in mixed stand.

|  | **Plant height** | **Tillers per plant** | **Grain number per tiller** | **Thousand grain weight** | **Grain weight per tiller** | **Harvest index** | **Heading date** |
| --- | --- | --- | --- | --- | --- | --- | --- |
| **Ppd-A1** | 19.5*** | 0 | 2 | 0.9 | 0.8 | 5.4* | 24.7*** |
| **Ppd-B1** | 0.1 | 2.7 | 0.1 | 1.8 | 0.6 | 0.1 | 15.7*** |
| **Ppd-D1** | 22.5*** | 3.3 | 0 | 3.1 | 0.6 | 2.9 | 10.5** |
| **Ppd-D1D2** | 21.0*** | 0.5 | 0 | 21.3*** | 2.1 | 0.4 | 6.1* |
| **VrnA1** | 7.1** | 0.5 | 0.2 | 0.1 | 0.1 | 0.2 | 2.8 |
| **Rht-B1** | 15.0*** | 0.4 | 1.3 | 7.1** | 4.5* | 0.2 | 2.7 |
| **Rht-D1** | 199.7*** | 2 | 4.5* | 0.2 | 2.4 | 21.8*** | 0.1 |
| **1B.1R** | 0.1 | 0.1 | 2.4 | 0 | 1.7 | 1.9 | 10.1** |
| **barc134-6B** | 24.2*** | 0.3 | 0 | 2.3 | 0.4 | 5.7* | 3.5 |
| **gwm165-4B** | 16.1*** | 1.9 | 0.1 | 30.6*** | 3.1 | 0.8 | 4.1* |
| **gwm165-4D** | 87.0*** | 0.3 | 4.1* | 5.3* | 0.1 | 5.0* | 1.1 |
| **gwm186-5A** | 1.8 | 0.5 | 3.5 | 0.4 | 5.4* | 1 | 6.9** |
| **gwm190-5D** | 1.1 | 0.1 | 0.6 | 0.2 | 0.6 | 0.3 | 2.3 |
| **gwm213-5B** | 5.4* | 6.8** | 5.5* | 13.4*** | 0.4 | 13.6*** | 0 |
| **gwm234-5B** | 1 | 0.1 | 0.5 | 0.1 | 0.2 | 3.2 | 2 |
| **gwm325-6D** | 6.5* | 2.7 | 2.1 | 1 | 2.5 | 1.8 | 0.1 |
| **gwm337-1D** | 34.2*** | 1.3 | 0.2 | 39.9*** | 4.4* | 3.5 | 0.5 |
| **gwm44-4A** | 8.4** | 2 | 4.9* | 0.1 | 2.6 | 0 | 1.3 |
| **gwm44-7D** | 2.2 | 1.4 | 0.3 | 6.7** | 2.6 | 0.2 | 4.2* |
| **gwm46-7B** | 2.3 | 3.8 | 1.5 | 10.3** | 0 | 1.3 | 1.9 |
| **gwm469-6D** | 0.6 | 0 | 0.3 | 10.5** | 2.9 | 4.7* | 0.1 |
| **gwm539-2D** | 40.8*** | 2.5 | 1.2 | 0 | 1.4 | 1.7 | 3.6 |
| **gwm583-5D** | 8.8** | 2.8 | 2.6 | 46.7*** | 1.1 | 0.1 | 2.2 |
| **gwm610-4A** | 1.8 | 10.2** | 0.2 | 0.5 | 0.4 | 0.2 | 2.4 |
| **gwm626-6B** | 6.8** | 3 | 1 | 4.5* | 0.1 | 2.1 | 23.0*** |
| **psp3100-1B** | 1.8 | 1.1 | 0 | 29.5*** | 3.8 | 1.4 | 7.4** |
| **psp3103A-4D** | 78.6*** | 1.1 | 0.7 | 1 | 0.9 | 3.4 | 0.6 |
| **wmc56_3B** | 13.8*** | 0.2 | 1.7 | 4.8* | 0.2 | 0.2 | 16.3*** |
| *,**,*** denotes P<0.05,P<0.01,and P<0.001, respectively, of the F-Test of marker main effect in the ANOVA | | | | | | | |

Table S 4. F-Test of the marker main effect in single genotype pure stand.

|  | **Plant height** | **Tillers per plant** | **Grain number per tiller** | **Thousand grain weight** | **Grain weight per tiller** | **Harvest index** | **Grain yield** |
| --- | --- | --- | --- | --- | --- | --- | --- |
| **Ppd-A1** | 4.8* | 0.2 | 1.3 | 0.1 | 0.8 | 3.4 | 0.1 |
| **Ppd-B1** | 1.2 | 1 | 2.1 | 0.9 | 1.1 | 0.3 | 1.5 |
| **Ppd-D1** | 0.1 | 12.0** | 0 | 0.8 | 0.6 | 0.5 | 1.2 |
| **Ppd-D1D2** | 0.3 | 0.4 | 0.5 | 0.2 | 1.5 | 0.2 | 1.5 |
| **VrnA1** | 2.1 | 0.4 | 0.8 | 1.3 | 3.9 | 1 | 1.7 |
| **Rht-B1** | 0.2 | 0.2 | 1.6 | 1 | 0.6 | 0.1 | 0.2 |
| **Rht-D1** | 9.0** | 3.6 | 15.5** | 2.2 | 6.2* | 9.0** | 8.2* |
| **1B.1R** | 1.3 | 2.9 | 0 | 0 | 0.1 | 0.2 | 0.1 |
| **barc134-6B** | 5.7* | 6.3* | 8.9** | 2 | 3.8 | 6.7* | 5.9* |
| **gwm165-4B** | 0.2 | 6.6* | 0.1 | 0.9 | 0.7 | 0.6 | 0.1 |
| **gwm165-4D** | 2.6 | 0 | 7.8* | 0 | 8.7** | 5.0* | 1.2 |
| **gwm186-5A** | 0.2 | 0.1 | 1.3 | 0.6 | 0.3 | 0.1 | 1.8 |
| **gwm190-5D** | 0.4 | 3.8 | 2.3 | 1.2 | 0.5 | 0.5 | 1.9 |
| **gwm213-5B** | 0.4 | 0.4 | 0.1 | 0 | 0.1 | 0.9 | 0 |
| **gwm234-5B** | 2.2 | 3.1 | 2.9 | 1.3 | 0.7 | 1.5 | 5.7* |
| **gwm325-6D** | 0.5 | 0.5 | 0 | 0 | 0 | 0 | 1.1 |
| **gwm337-1D** | 0.7 | 0.3 | 1.2 | 1.8 | 0.1 | 1.5 | 0 |
| **gwm44-4A** | 0.4 | 1.2 | 2.4 | 0.7 | 0.9 | 1.3 | 4.8* |
| **gwm44-7D** | 0.2 | 0.2 | 0.3 | 0 | 0.1 | 0.2 | 0 |
| **gwm46-7B** | 0.3 | 2.2 | 0 | 0.5 | 0.3 | 0.1 | 0.1 |
| **gwm469-6D** | 0.4 | 7.8* | 0.4 | 3.4 | 4.8* | 1 | 0 |
| **gwm539-2D** | 0.3 | 1.4 | 2.5 | 2.8 | 0.2 | 2.6 | 4 |
| **gwm583-5D** | 0 | 0.1 | 0.7 | 0.1 | 0.6 | 0.4 | 0 |
| **gwm610-4A** | 1.2 | 0 | 0.6 | 0.6 | 2.6 | 0.1 | 0.2 |
| **gwm626-6B** | 0.3 | 5.4* | 1.2 | 3.5 | 0 | 0 | 2.8 |
| **psp3100-1B** | 0.3 | 0 | 0.5 | 0.4 | 1.6 | 0.1 | 0.5 |
| **psp3103A-4D** | 1.5 | 0.4 | 2 | 0.2 | 1.4 | 1.1 | 2.3 |
| **wmc56_3B** | 0.9 | 4.6* | 0.1 | 0 | 0 | 0.6 | 2.8 |
| *,**,*** denotes P<0.05,P<0.01,and P<0.001, respectively, of the F-Test of marker main effect in the ANOVA | | | | | | | |

***Supporting Figures***

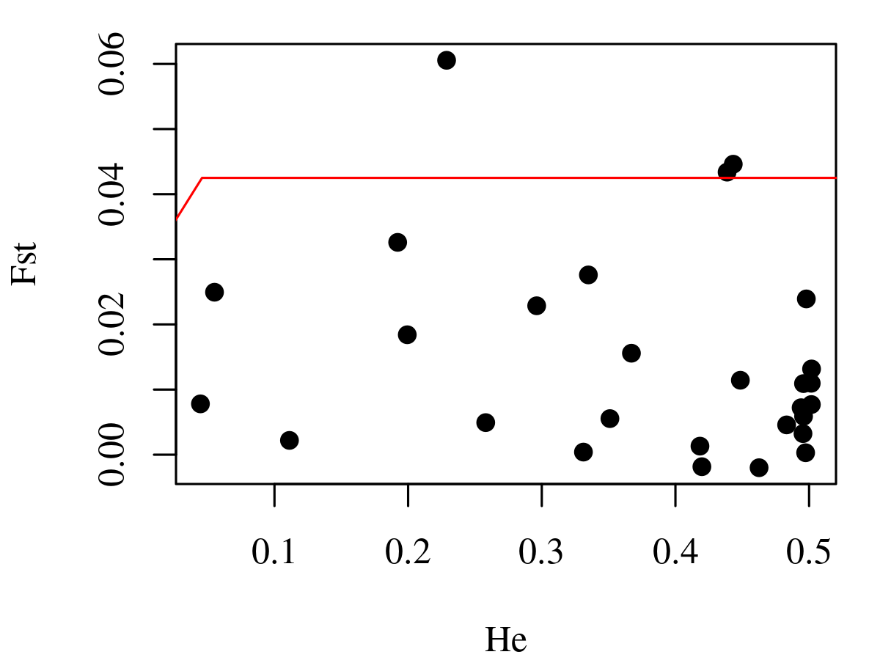

Fig. S1 F_st_ values at generation 10 comparing all 4 bread wheat CCP populations plotted against expected heterozygosity (H_e_). The red line indicates the 95% null distribution quantile from the island model simulation with *Arlequin* based on 20,000 simulations with 100 demes (see Material and methods). The three dots above the red line represent the three loci identified as under selection (P<0.05).

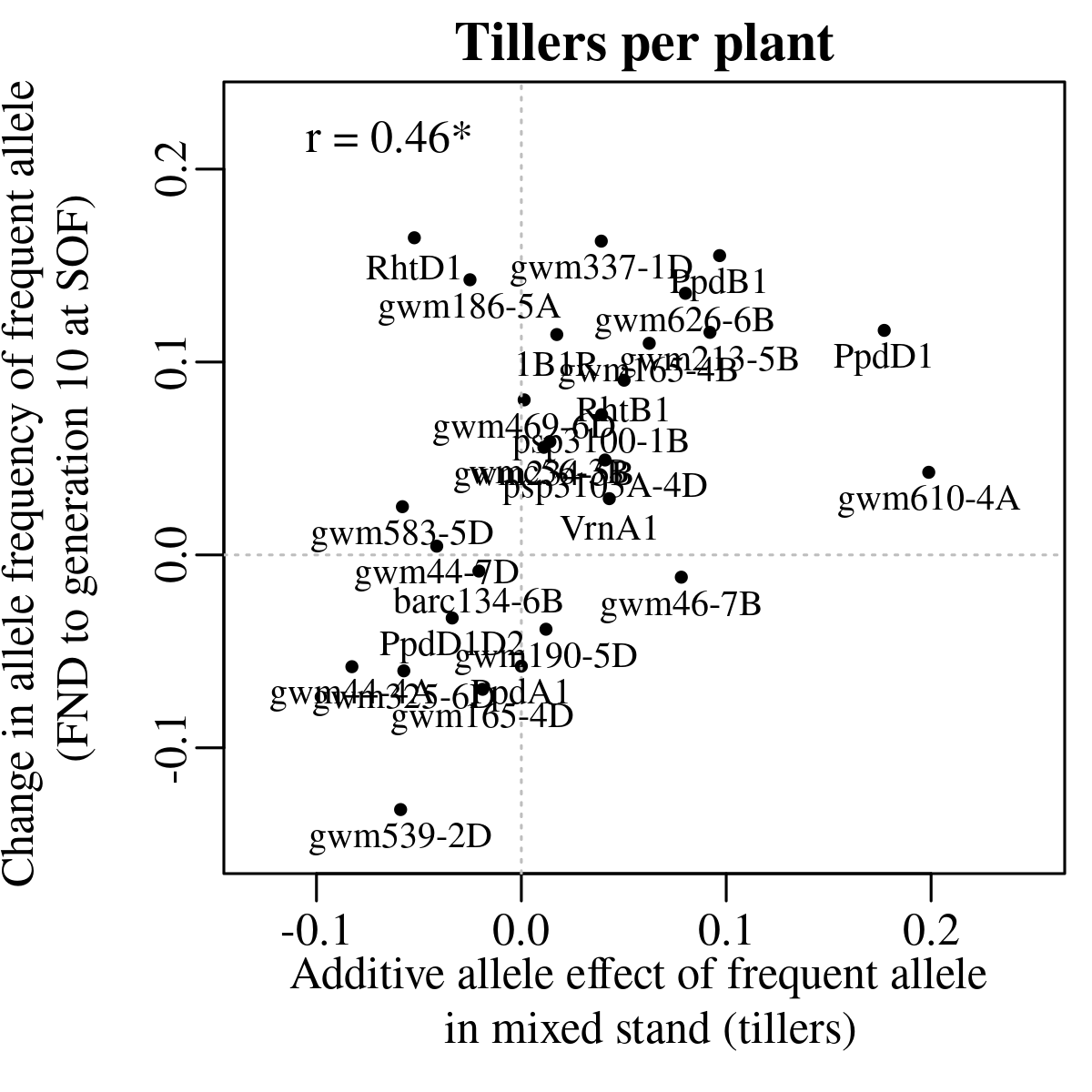

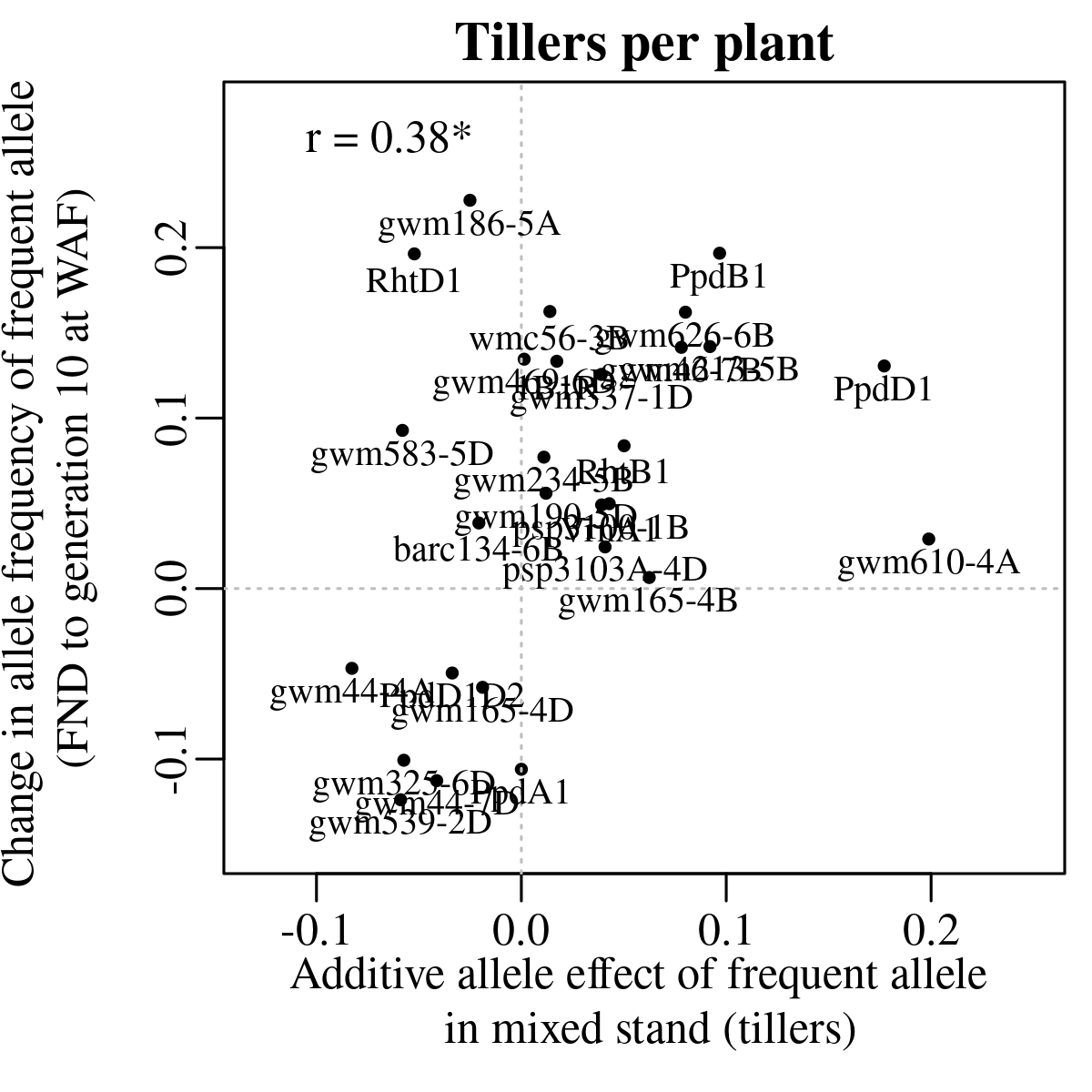

Fig. S2. Relation of the additive allele effect on tillers per plant in single plants in mixed stands to the change in allele frequency at SOF (left) and WAF (right).
